# Controlling bias and inflation in epigenome- and transcriptome-wide association studies using the empirical null distribution

**DOI:** 10.1101/055772

**Authors:** Maarten van Iterson, Erik van Zwet, the BIOS Consortium, P. Eline Slagboom, Bastiaan T. Heijmans

**Author notes:** Correspondence to: M. van Iterson, Leiden University Medical Center, Department of Molecular Epidemiology, Leiden, The Netherlands.

## Abstract

Association studies on omic-level data other then genotypes (GWAS) are becoming increasingly common, i.e., epigenome-and transcriptome-wide association studies (EWAS/TWAS). However, a tool box for the analysis of EWAS and TWAS studies is largely lacking and often approaches from GWAS are applied despite the fact that epigenome and transcriptome data have vedifferent characteristics than genotypes. Here, we show that EWASs and TWASs are prone not only to significant inflation but also bias of the test statistics and that these are not properly addressed by GWAS-based methodology (i.e. genomic control) and state-of-the-art approaches to control for unmeasured confounding (i.e. RUV, sva and cate). We developed a novel approach that is based on the estimation of the empirical null distribution using Bayesian statistics. Using simulation studies and empirical data, we demonstrate that our approach maximizes power while properly controlling the false positive rate. Finally, we illustrate the utility of our method in the application of meta-analysis by performing EWASs and TWASs on age and smoking which highlighted an overlap in differential methylation and expression of associated genes. An implementation of our new method is available from http://bioconductor.org/packages/bacon/.

---

The large-scale analysis of epigenome and transcriptome data in population studies is thought to answer fundamental questions about genome biology and will be instrumental in linking genetic and environmental influences to disease etiology^1^. Worldwide, research groups are now joining forces to generate and analyze such data^2–6^ complementary to the vast resources of genetic data that are already present and have been successfully used in Genome-Wide Association Studies (GWASs). While the analysis tool box for GWAS has matured, the development of effective methodology for the analysis of epigenome-and transcriptome-wide association studies (EWAS and TWAS) is a nascent field of research. In an EWAS, DNA methylation levels of typically 100 thousands of CpG dinucleotides are individually tested for an association with an outcome of interest, while in a TWAS this is done for expression levels of 10 thousands of genes. Currently, the analysis of EWASs and TWASs heavily relies on approaches specifically designed for GWAS^7^. However, epigenome and transcriptome data are crucially different from genetic data. They are quantitative measures, and not discrete like genotypes, that are subject to major effects of technical batches and biological influences, including cellular heterogeneity, prone to introduce confounding. Moreover, the associations between complex diseases and their risk factors often are more widespread for epigenome and transcriptome data than for genetic data.

A key aspect in the analysis of ome-wide association studies is addressing test-statistic inflation, which leads to an overestimation of the level of statistical significance and can dramatically increase the number of false positive findings^8^. This has always been a major concern in GWAS, but is also observed in EWAS^9,10^ and often the level of inflation exceeds that observed in GWAS. In GWAS, test statistic inflation is commonly addressed using genomic control in which the test statistics are divided by the genomic inflation factor (λ_GC_) that estimates the inflation by comparing observed test statistics across all genetic variants evaluated to those expected by chance^9,10^. Recent work pointed out crucial limitations of genomic control in GWAS^11,12^. Notably, genomic control was shown to provide an invalid estimate of test statistic inflation when the outcome of interest is associated with many, small genetic effects^11^. In EWAS and TWAS, this is the rule rather than exception. Moreover, it has been shown that test statistics may not only be subject to inflation but also to bias^13^, which is not corrected for when using genomic control. Bias leads to a shift in the distribution of effect-sizes and is driven by confounding, a prominent feature of EWAS and TWAS but not GWAS. Together, this calls for the development of new methods specifically designed to address inflation and bias in EWAS and TWAS analyses.

Although generally ignored, the fact that genomic control results in an overestimation of the actual inflation unless it is estimated on the basis of genetic variants not associated with the outcome of interest was originally noted^8,14^ (**Box 1**). As a solution, it was proposed to estimate λ_GC_ assuming a fixed but small number of associated genetic variants (i.e. 10) using a Bayesian outlier model^15^. Although an improvement for GWAS with few associations, it will not be sufficient to solve the overestimation of inflation in EWAS and TWAS that typically yield substantially more associations, nor does it address the occurrence of bias. In other fields, alternative approaches have been proposed for large-scale multiple testing problems with invalid null distributions by using empirical null distributions that obviate inflation and bias^16–19^. The utility of these approaches in EWAS and TWAS, however, remains to be evaluated.

Here, we used simulations and large methylome (n=2203) and trancriptome (n=1910) data^20,21^ to show that correcting test statistic inflation using genomic control is too conservative for EWAS and TWAS and that bias in test statistics cannot be ignored. Moreover, we demonstrate that the bias and inflation in fact are the mean and variance estimates of the empirical null distribution, which can be estimated with a Bayesian algorithm. Application of state-of-the-art batch correction methods, like, ruv, sva and cate^22–24^, are not able to completely remove all bias and inflation and the resulting test-statistics require calibration to achieve optimal statistical power while controlling the number of false positives at the desired level. We present a fast and user-friendly software implementation of this new method called bacon. Finally, we show the utility of our approach by performing an EWAS and TWAS meta-analysis of two commonly studied outcomes, namely age and smoking status.

## RESULTS

### The genomic inflation factor is not suitable to measure inflation in EWAS/TWAS

We performed an EWAS and TWAS of age and smoking status using a subset of two population cohorts containing 500 individuals each (**Supplementary Table 1**). The analyses were adjusted for known biological (including measured white blood cell counts) and technical covariates within a linear model framework (**Supplemental Methods**). Test-statistic inflation was observed for both cohorts, data types and outcomes (**Fig. 1**). The amount of inflation, quantified using the λ_GC_^8^ (Box 1), ranged between 1.33–1.72 for the EWASs and between 1.21–1.54 for the TWASs (**Table 1**). The levels of inflation appeared to be outcome-specific: the inflation was consistently higher for age than smoking status (**Fig. 1** and **Table 1**). Also, there was a cohort-specific effect: the inflation for the age EWAS was much higher for the cohort, LifeLines, with a wide age range (34-55 years; λ_GC_=1.72) than for the cohort, Leiden Longevity Study, with the narrow age range (55–64 years; λ_GC_=1.52). The level of inflation appeared to be correlated with the expected amount of true associations^2–6^. For example, for smoking status the expression of a moderate number of genes and methylation levels CpG dinucleotides are known to be associated^5,6^, while the number of associations present for age is much greater^2–4^. A simulation study substantiated this impression (**Fig. 2** and **Supplemental Methods**) and the dependence of λ_GC_ on the number of true associations could be shown mathematically (**Box 1** and **Supplemental Text**) in line with previous work that reported this relationship for the Armitage trend test^8,14^. We conclude that in EWASs and TWASs the inflation of test statistics is commonly overestimated when using λ_GC_.

**Figure 1.**
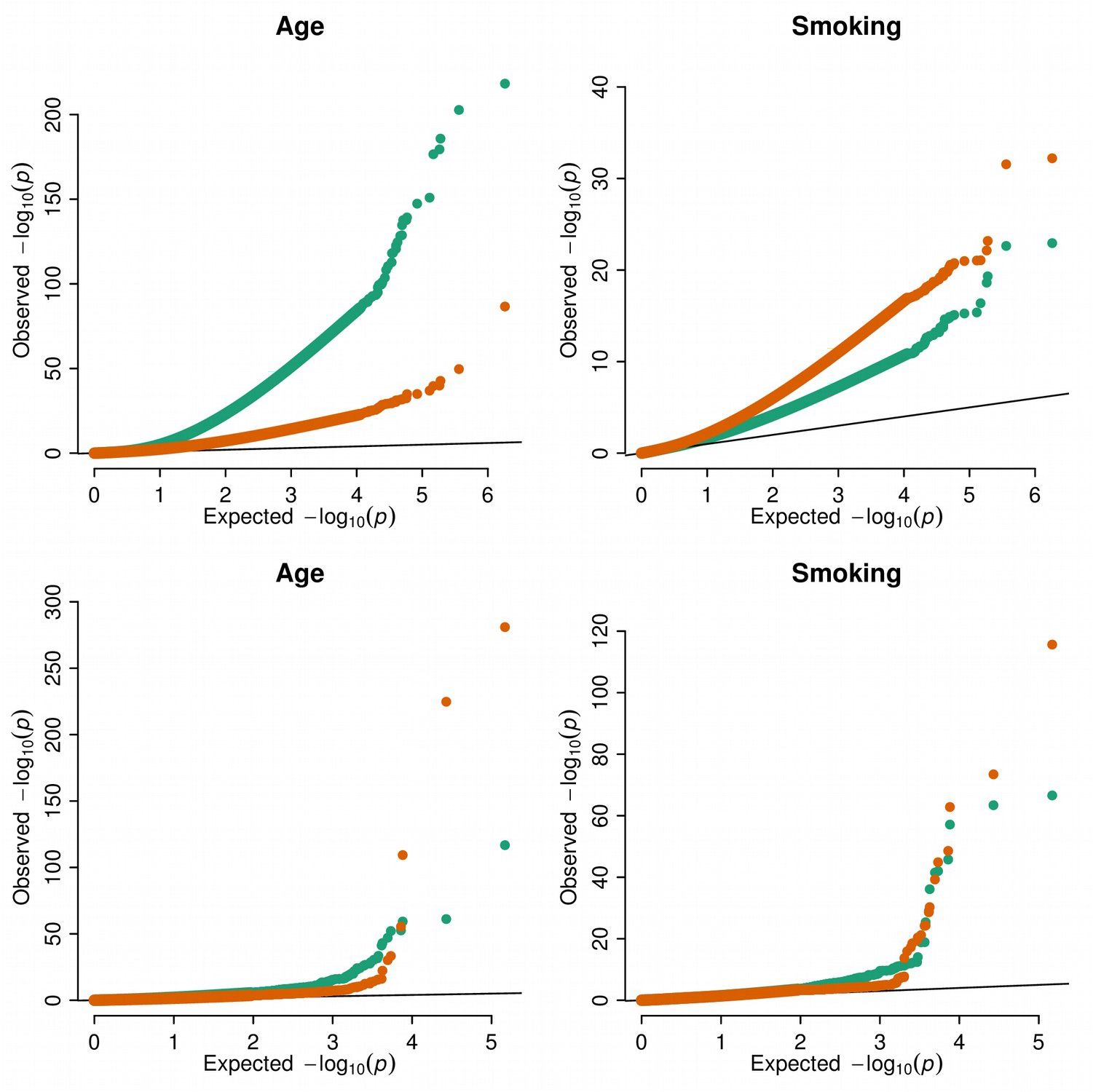
Inflated epigenome-and transcriptome-wide association studies. Quantile-quantile plots for EWASs (**a** and **b**) and TWASs (**c** and **d**) on two cohorts LifeLines and Leiden Longevity Study for phenotypes age and smoking status. QQ-plots shows the observed log10-transformed P values obtain from a linear model corrected for known biological-and technical covariates against quantiles from the theoretical null distribution. Strong inflation is observed for both the EWAS and TWAS for age while for smoking status the inflation is much smaller.

**Table 1.**
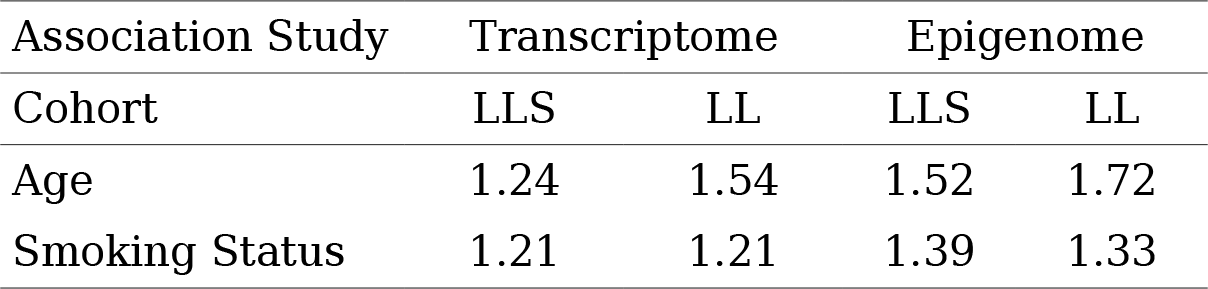
Genomic inflation factors for the EWAS and TWAS shown in Figure 1. Inflation factors are calculated as the square-root of the median of squared test-statistics divided by 0.456, the median of chi-square distribution with one degree of freedom^8^.

| Association Study | Transcriptome |  | Epigenome |  |
| --- | --- | --- | --- | --- |
| Cohort | LLS | LL | LLS | LL |
| Age | 1.24 | 1.54 | 1.52 | 1.72 |
| Smoking Status | 1.21 | 1.21 | 1.39 | 1.33 |

**Figure 2.**
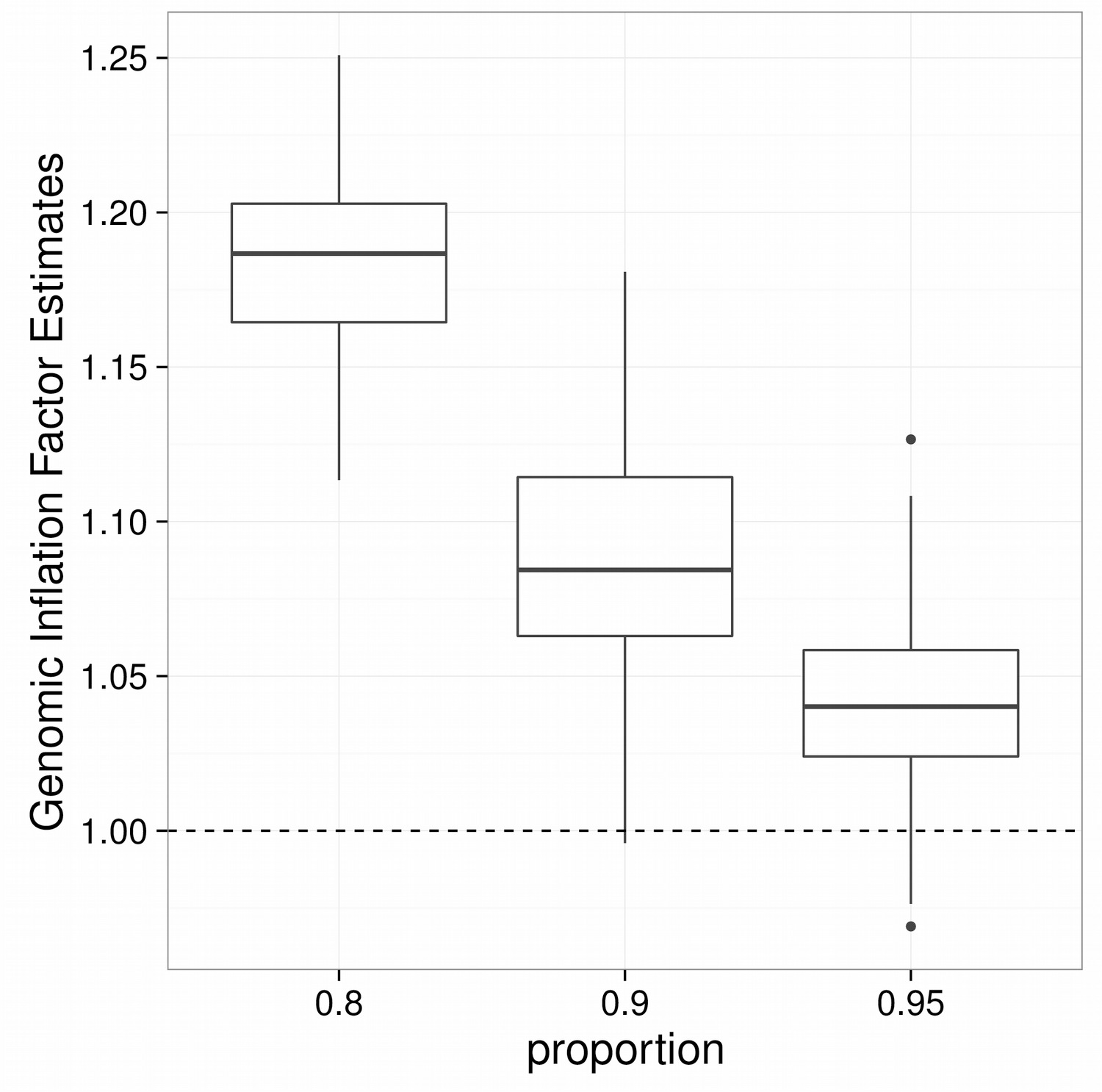
The genomic inflation factor overestimates inflation if a moderated proportion of true associations is present. Sets of test-statistics were generated with differentv amounts of true associations (20%, 10% and 5%) but without any true inflation, i.e., the inflation factor should be equal to one (**Supplemental Methods**). The genomic inflation factor was calculated as the square-root of the median of squared test-statistics di ided by 0.456, the median of chi-square distribution with one degree of freedom^8^.

### EWAS/TWAS not only suffer from test-statistics inflation but biases too

While quantile-quantile plots of expected versus observed test statistics, or their corresponding *P* values, are frequently used to visualize test-statistic inflation (**Fig. 1**), the alternative representation through a histogram of the test-statistics reveals a second artifact namely a bias in the test-statistics (**Fig. 3a** and **Supplementary Fig. 2**). This bias is visible as a deviation of the mode of the observed statistics from zero; the mode of the standard normal distribution. Since the majority of features, being genetic variants, CpGs or genes, will not be associated with the outcome of interest, statistical theory assumes that test-statistics obtained from a linear model follows a standard normal distribution that is centered at zero. We observed bias in the EWASs and TWASs of age and smoking irrespective of cohort and outcome (**Supplementary Fig. 2**). Genomic control does not address bias because it uses a normal distribution with the mode fixed at zero (**Box 2** and **Supplemental Text**). The misspecification of the observed distribution of test statistics by genomic control is illustrated in **Figure 2c**. Of note, also permutation-based approaches, which are often assumed to rescue violations of assumptions regarding the theoretical null distribution, do not result in a proper null distribution and bias and inflation remains^16,25^ (**Fig. 2d** and **Supplemental Text**).

**Figure 3.**
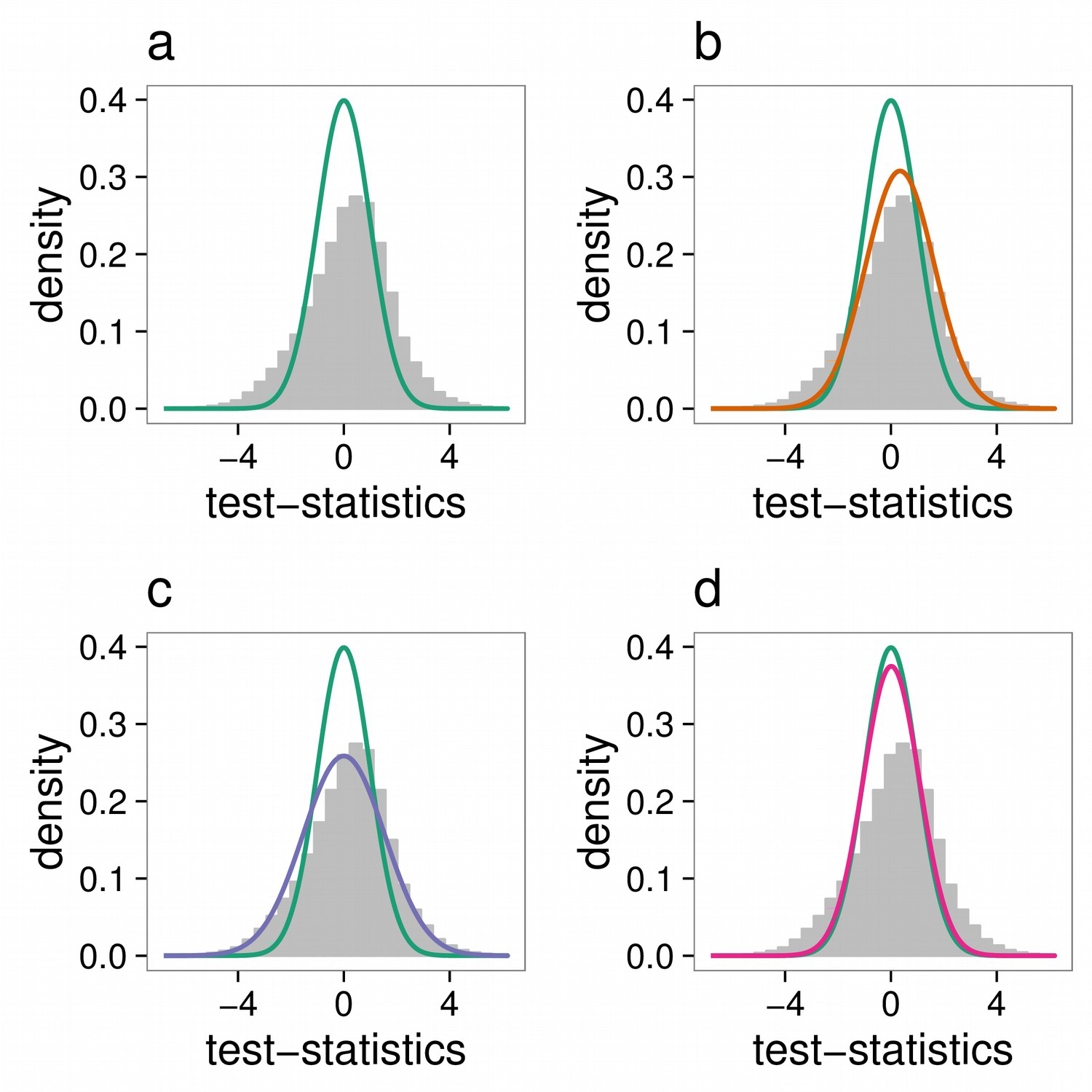
Bias in transcriptome-wide association studies. Histogram of test-statistics from the TWAS on age for the LifeLines cohort. Each panel a different null distribution is shown. **a)** a standard normal distribution with mean and variance (0.0, 1.0), **b)** the empirical null; normal distribution with estimated mean and variance using our novel Bayesian approach (0.23, 1.5^2) **c)** normal distribution with zero mean and variance equal to the estimate genomic inflation factor (0.0, 1.5^2) and **d)** normal distribution with permutation based estimates of mean and variance (−0.006, 1.1^2).

### Estimating bias and inflation

To detect bias and inflation in EWAS and TWAS, we developed a method that estimates the empirical null distribution of test statistics using a Bayesian statistical approach. The method fits a three component normal mixture to the observed distribution of test-statistics using a Gibbs Sampling algorithm^26^. One component is forced to represent the null distribution with mean and standard deviation representing the bias and inflation. The other two components, one smaller than zero the other larger, capture the fraction of true associations present in the data, which is assumed to be an unknown minority of tests (**Supplemental Methods**). Hence, our method simultaneously provides parametric estimates for the distribution of test-statistics not associated with the outcome of interest (i.e. the empirical null distribution), and adapts to test-statistics bias without being affected by an unknown proportion of true associations (**Fig. 2b** and **Supplementary Fig. 2**). We compared our method to derive the empirical null distribution to previously proposed methods^16^ in a simulation study. This showed that the performance of our method is equal or better than other methods under different scenarios. Importantly, our method resulted in the most stable estimation of the inflation which suggests that other methods randomly over-or underestimate the level of bias and inflation (**Supplemental Methods** and **Supplementary Fig. 3)**.

### Correction for unobserved covariates reduces test-statistic bias and inflation

The primary causes of inflation and bias are thought to be unmeasured technical and biological confounding^16,27^, e.g., population-substructure or batch effects. Various methods have been developed to control for these factors in high-dimensional data^22–24, 28–31^. We applied four methods to adjust an EWAS and TWAS of age in one cohort of 500 individuals and investigated their impact on bias and inflation. All approaches reduced the bias and inflation as compared with a model using known covariates only (**Table 2**, **Supplemental Methods** and **Supplementary Table 2**). Nevertheless, residual bias and inflation was observed. Therefore, we designed a two-stage approach in order to preserve statistical power while appropriately controlling the number of false positives. First, we performed an analysis that correct for known biological and technical covariates plus estimated unobserved covariates, followed by estimating and adjusting the residual bias and inflation using the empirical null distribution. In the adjustment step, *P* values are calculated using the empirical null distribution instead of the standard normal or overdispersed or inflated normal that is used by the genomic control approach. It is crucial to note, that bias not only results in incorrect test statistics and *P* values but also results in biased effect size estimates (**Box 2** and **Supplemental Methods**).

To evaluate the performance of the two-stage approach, we conducted a numerical simulation. To account for unmeasured confounding, we selected cate, a state-of-the-art method that was shown to have superior performance in estimating unobserved covariates as compared with alternative approaches^24^. Under three different scenarios, our method in combination with cate yielded the highest power with the fraction of false positives close to the nominal level (0.058±0.0052, 0.059±0.0055 and 0.059±0.0059), whereas approaches that ignore unobserved covariates lead to high false positive rates and those that use genomic control resulted in lower power (**Table 3** and **Supplemental Methods**). Also test-statistic calibration that was proposed to use in combination with cate^24^ and is closely related to genomic control, was conservative resulting in lower power.

**Table 2.**
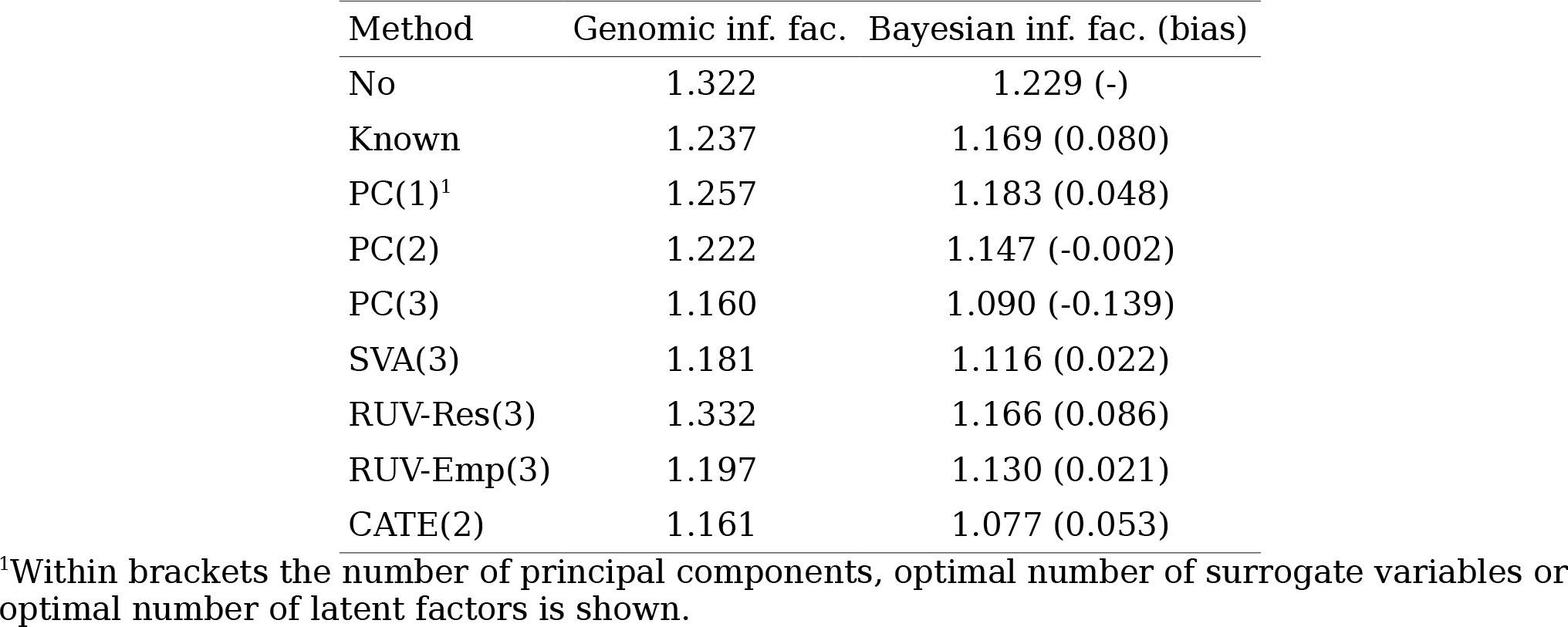
Correction for unknown batches reduces the inflation in a TWAS on age. Genomic and Bayesian inflation factors (and biases) calculated from test-statistics obtained by fitting linear models with 1) only known covariates 2, 3, and 4) known covariates plus one, two and three principal component 5) plus three optimal surrogate variables estimated using SVA^22^ 6) plus three unobserved covariates estimated using RUV^29^ with the residual-method 7) plus three unobserved covariates estimated using RUV^29^ with the empirical-method and 8) plus two optimal latent variables estimated using CATE^24^.

| Method | Genomic inf. fac. | Bayesian inf. fac. (bias) |
| --- | --- | --- |
| No | 1.322 | 1.229 (-) |
| Known | 1.237 | 1.169 (0.080) |
| PC(1)<sup>1</sup> | 1.257 | 1.183 (0.048) |
| PC(2) | 1.222 | 1.147 (-0.002) |
| PC(3) | 1.160 | 1.090 (-0.139) |
| SVA(3) | 1.181 | 1.116 (0.022) |
| RUV-Res(3) | 1.332 | 1.166 (0.086) |
| RUV-Emp(3) | 1.197 | 1.130 (0.021) |
| CATE(2) | 1.161 | 1.077 (0.053) |

<sup>1</sup>Within brackets the number of principal components, optimal number of surrogate variables or optimal number of latent factors is shown.

**Table 3.** Numerical simulation with unobserved covariates. Mean and standard deviation of Type I error and Power for 100 simulations using different approaches to reduced the impact of unobserved covariates for three different scenario’s. A scenario with 5 or 10 unobserved covariates and 10% true associations or 5 unobserved covariates with 15% true associations. Number of samples and number of features in each scenario was set to 100 and 2000, respectively. The different approach are naÏve) a linear model ignoring unobserved covariates, naÏve-gc) naÏve with genomic control, naÏve-bc) naÏve with control of bias and inflation using bacon, cate) cate.calibrate) cate.bacon and oracle) linear model with the simulated unobserved covariates.

|  | 5 unobs. cov./prop 0.9 |  | 10 unobs. cov./prop. 0.9 |  | 5 unobs. cov./prop 0.85 |  |
| --- | --- | --- | --- | --- | --- | --- |
|  | Type I error | Power | Type I error | Power | Type I error | Power |
| naive | 0.72 (0.036) | 0.72 (0.049) | 0.58 (0.043) | 0.59 (0.055) | 0.71 (0.037) | 0.71 (0.043) |
| naive-gc | 0.00091 (0.0021) | 0.0048 (0.0073) | 0.023 (0.0083) | 0.028 (0.013) | 0.0014 (0.0029) | 0.0048 (0.0054) |
| naive-bc | 0.029 (0.0076) | 0.047 (0.018) | 0.045 (0.0054) | 0.053 (0.016) | 0.029 (0.0073) | 0.045 (0.014) |
| cate | 0.06 (0.0056) | 0.86 (0.037) | 0.069 (0.0061) | 0.85 (0.042) | 0.062 (0.0059) | 0.86 (0.036) |
| cate.calibrate | 0.03 (0.0042) | 0.77 (0.053) | 0.032 (0.0053) | 0.74 (0.055) | 0.021 (0.0046) | 0.7 (0.057) |
| cate.bacon | 0.058 (0.0052) | 0.86 (0.041) | 0.059 (0.0055) | 0.83 (0.047) | 0.059 (0.0059) | 0.86 (0.04) |
| oracle | 0.052 (0.0052) | 0.85 (0.039) | 0.053 (0.0051) | 0.83 (0.044) | 0.053 (0.0053) | 0.85 (0.037) |

### Bayesian control for fixed-effect meta-analysis

A main development in the field of EWAS and TWAS, analogous to current practice in GWAS, is the combined analysis of multiple population studies to detect an increasing number of associations including those with small effect-sizes. Fixed-effect meta-analysis combines estimated effect-sizes and their standard errors from different studies to construct pooled estimates resulting in higher precision and hence superior statistical power^32^.

We performed an EWAS and TWAS of age and smoking status in four cohorts totaling 2203 and 1910 samples, respectively. We combined the results through fixed-effect meta-analysis after correction for bias and inflation in the individual cohorts (**Fig. 4** and **Table 4**). The level of bias and inflation remained present despite addressing unmeasured confounding using cate. Of note, estimates of inflation using genomic control were much more variable across analyses and cohorts and adjustment of test statistics using λ_GC_ can be expected to result in invalid results for a subset of cohorts. Our method fully removed all bias and inflation. Critically, bias (<0.03) and inflation (<1.14) remained minimal in the meta-analysis compared to an meta-analysis using genomic control (**Table 4**). The top-hits identified for age and smoking included those consistently reported^2–6^. Furthermore, our simultaneous performance of an EWAS and TWAS in a large meta-analysis showed a remarkable overlap in results, 410 and 57 unique genes for the association studies on age and smoking status, respectively (assigning the nearest gene to a CpG probe) (**Supplementary Table 3a, 3b, 3c** and **3d**). For example, both DNA methylation near and expression of *CD248, DNMT3A* and *FBLN2* was associated with age (**Fig. 5a**), while the same was true for *GPR15, AHRR* and *CLDND1* for smoking (**Fig. 5b**). In total 15967 (3.5%) CpG loci and 1020 (2.7%) genes were significantly associated with age (Bonferroni corrected *P* values <0.05) for smoking status 1128 (0.25%) CpG loci and 301 (0.80%) genes.

**Figure 4.**
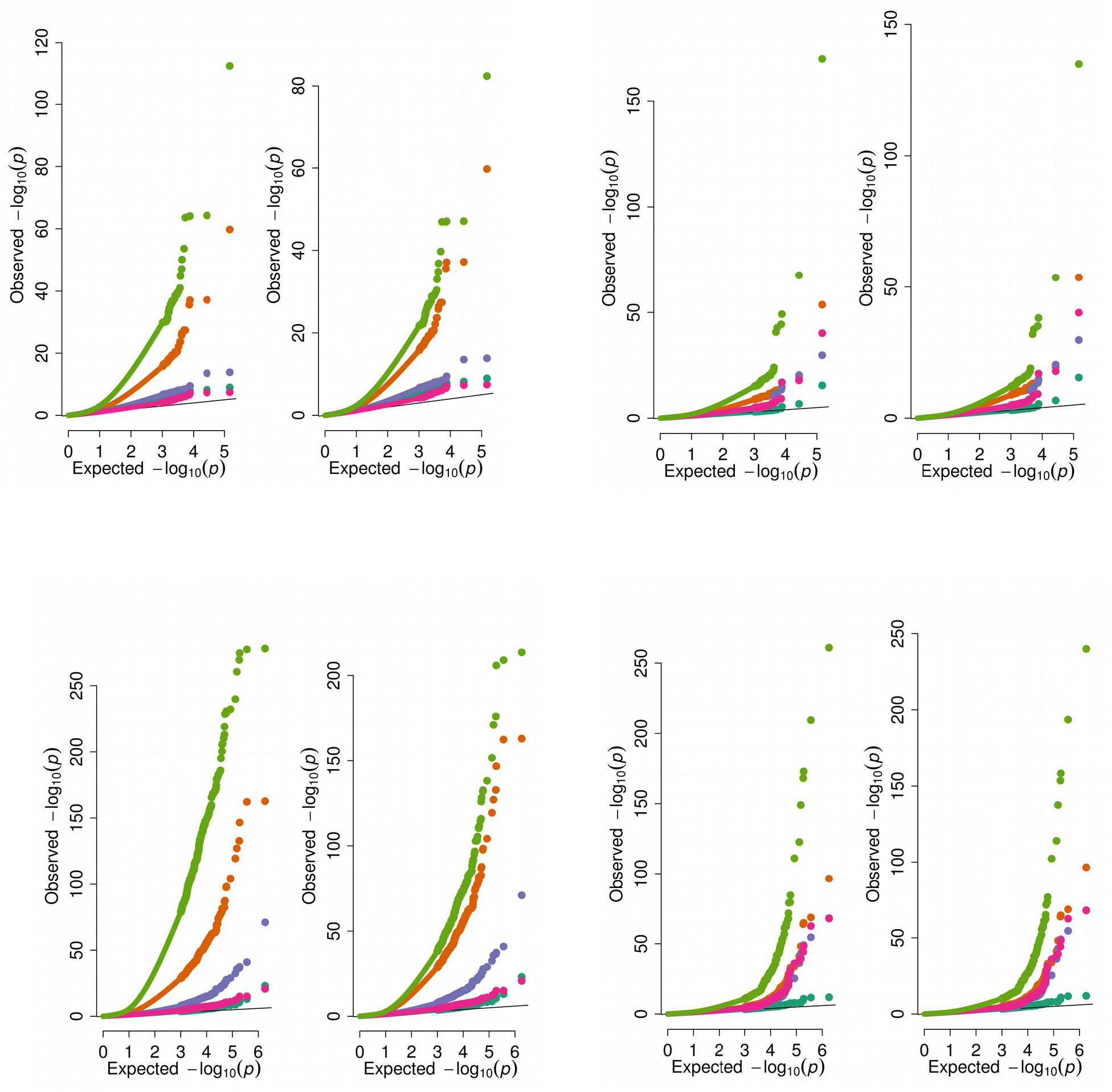
Bayesian control for fixed-effect meta-analyses-. Quantile-quantile plots for meta-analysis for TWAS (**a** and **b**) and EWAS (**c** and **d**) on Age (**a** and **c**) and Smoking Status (**b** and **d**). In each plot left-panels represents bias and inflation uncorrected qq-plots and right-panels the bias and inflation corrected qq-plots.

**Table 4.**
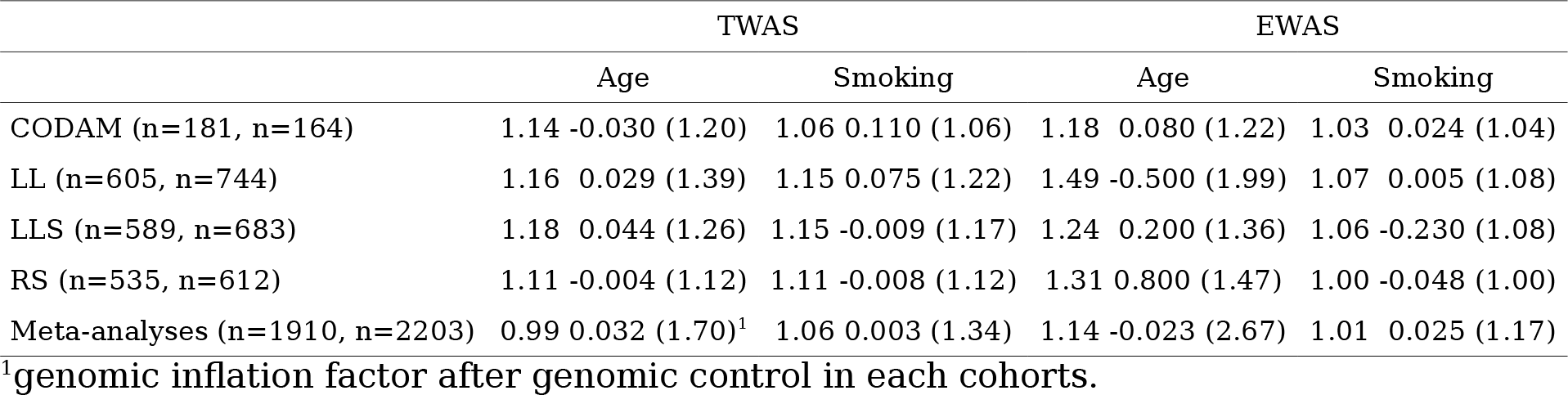
Inflation factors for the different EWAS/TWAS that were part of the meta-analyses. EWAS/TWAS meta-analysis conducted on four cohorts. Table represents the Bayesian inflation/bias and genomic inflation factor within brackets for each cohort and phenotype tested. In all analyses we used CATE to estimated three unobserved covariates, which were added to the linear model.

|  | TWAS |  | EWAS |  |
| --- | --- | --- | --- | --- |
|  | Age | Smoking | Age | Smoking |
| CODAM (n=181, n=164) | 1.14 -0.030 (1.20) | 1.06 0.110 (1.06) | 1.18 0.080 (1.22) | 1.03 0.024 (1.04) |
| LL (n=605, n=744) | 1.16 0.029 (1.39) | 1.15 0.075 (1.22) | 1.49 -0.500 (1.99) | 1.07 0.005 (1.08) |
| LLS (n=589, n=683) | 1.18 0.044 (1.26) | 1.15 -0.009 (1.17) | 1.24 0.200 (1.36) | 1.06 -0.230 (1.08) |
| RS (n=535, n=612) | 1.11 -0.004 (1.12) | 1.11 -0.008 (1.12) | 1.31 0.800 (1.47) | 1.00 -0.048 (1.00) |
| Meta-analyses (n=1910, n=2203) | 0.99 0.032 (1.70)<sup>1</sup> | 1.06 0.003 (1.34) | 1.14 -0.023 (2.67) | 1.01 0.025 (1.17) |

<sup>1</sup>genomic inflation factor after genomic control in each cohorts.

**Figure 5.**
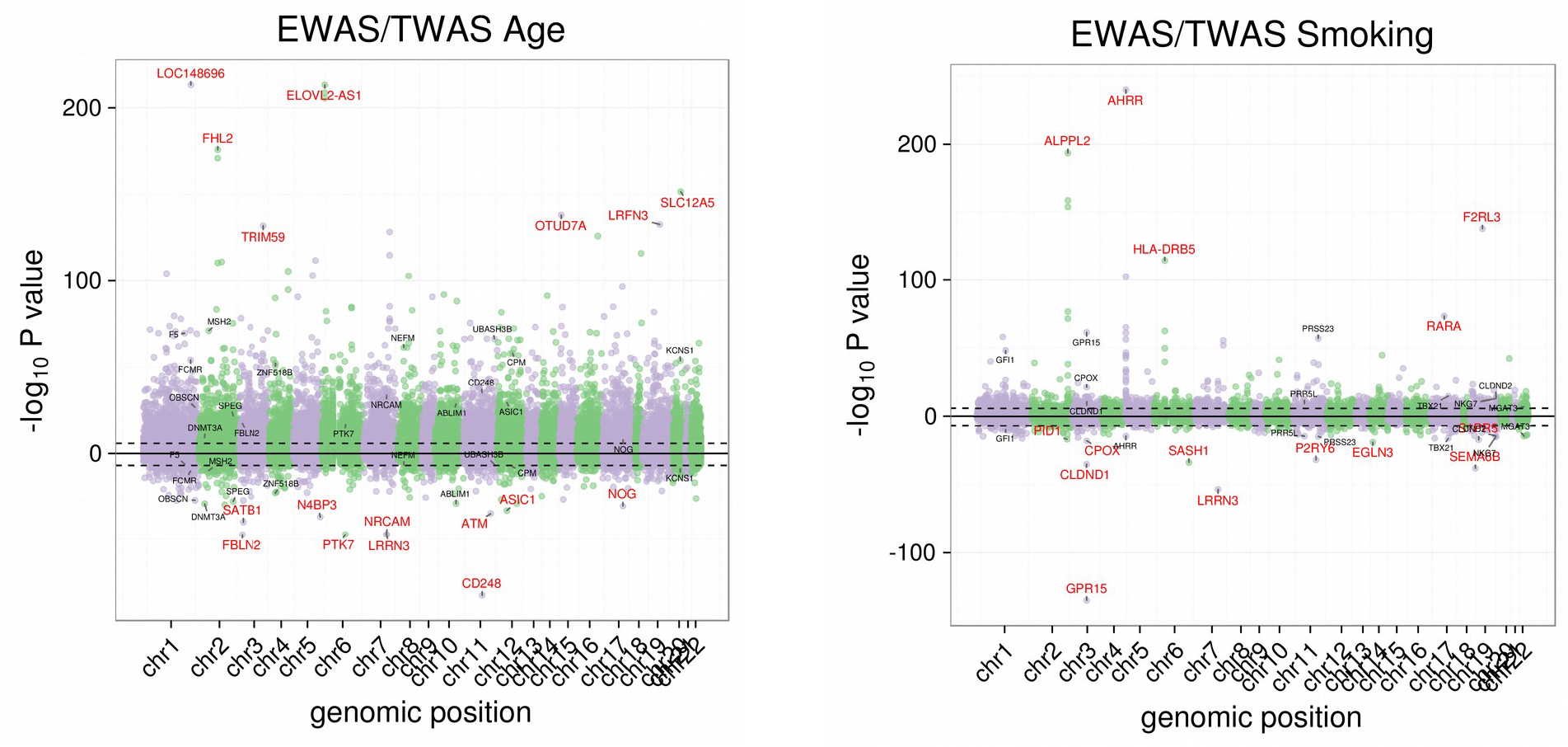
Manhattan plots for meta-analyses results for EWAS and TWAS of age and smoking-. Panel a show the −log10 P values from the meta-analysis of age were the positive part of the y-axis show the results for the EWAS while the negative part of the TWAS. Black dashed lines indicated 0.05 Bonferroni thresholds. In red are highlighted the top 10 genes which for the EWAS are the genes closest to the significant CpG locus. In black are highlighted the top 10 genes that are both significant for the EWAS as well as the TWAS.

## DISCUSSION

We describe a novel Bayesian approach to detect and correct bias and inflation in EWASs and TWASs which has the crucial characteristic that it is largely independent of the fraction of true associations in the data. We showed that genomic control, that is commonly used in GWAS, results in spurious association because it does not address bias and, moreover, inflation adjustment on the basis of genomic control reduces power because it increases with the fraction of true associations. We implemented our method in the software package bacon including bias and inflation corrected fixed-effect meta-analyses that are increasingly being used. Moreover, the performance of our approach towards estimating the empirical null distribution of test statistics performs better or as good as existing approaches^16^ by taking advantage of prior knowledge of the distribution and the composition of test-statistics.

Methods that try to estimate unmeasured covariates^22–24^ and those that try to recover the empirical null distribution are based on same principle: extracting information from features that are not associated with the outcome of interest to control the impact of unobserved covariates. For example, RUV^23^, extracts this information from control probes that, *a priori,* should not be associated with the phenotype of interest but can reflect the presence of unobserved covariates or unwanted variation. Recently, Wang *et al.*^24^ showed that such state-of-the-art approaches^22, 23^ can be unified in a single mathematical framework, which was implemented in cate. Genomic control, as proposed by Devlin and Roeder^8^, is based on estimating an inflation factor from features known not to be associated with the phenotype of interest but in practice are estimated using all data because those unassociated features are not known. Existing methods that estimate the complete empirical null distribution similarly assume that majority of features are not associated^16–19^. Although our method bacon is related to the latter approaches, it is designed to be much more flexible in dealing with larger fractions of true associations, which turns out to be crucial in particular for EWAS and TWAS meta-analyses.

In the present paper we describe how to correct for bias and inflation remaining after state-of-the-art control for unmeasured confounding factors. Our study cannot identify the causes of the remaining bias and inflation. In general, the potential causes include failed mathematical assumptions, correlation across sampling units, correlation across cases and the presence of unobserved covariates^16^. While we have attributed the latter cause in detail, the first cause, of failed mathematical assumptions, is of particular relevance for EWAS/TWAS. Since, most EWAS/TWAS use a linear regression approach, whereas a Poisson or negative binomial regression model might be more appropriate for a TWAS using RNA-seq count data^33,34^. Here we have chosen to use appropriate data transformations that allows us to use the linear model framework for association analyses^35,36^.

Controlling bias and inflation in EWAS/TWAS and meta-analyses thereof is critical to obtain correct inference. Often the high-dimensionality of omics data is seen as a burden for the statistical analyses. However, statistical approaches that are based on empirical Bayes methods turn the dimensionality into advantage. Our method bacon is such a method. The fact that it is *a priori* known that the majority of features will not be associated (without the requirement for specific assumptions as to the precise fraction) facilitates the derivation of an empirical null distribution that can be applied to achieve a correct inference, even in situations were unmeasured confounding remains undetected or is subject to measurement error.

Our work extends the work of Devlin and Roeder^8^, who originally propose to use genomic control to tackle test-statistic inflation for GWAS and links their method to the pioneering work of Efron^16^ on estimating an empirical null distribution for high-dimensional data inference. Hence, although specifically applied to EWAS and TWAS, our statistical approach may have implications for any field focusing on inference for high-dimensional data.

## AUTHORS CONTRIBUTIONS

MvI and BTH designed the study. MvI and EvZ developed the statistical method. MvI performed all statistical analyses and implemented the software. MvI and BTH wrote the paper. PS revised manuscript critically for important intellectual content. All authors read and approved the manuscript.

## Competing financial interests

The authors declare no competing financial interests.

## Acknowledgement

This work was done within the framework of the Biobank-Based Integrative Omics Studies (BIOS) Consortium funded by BBMRI-NL, a research infrastructure financed by the Dutch government (NWO 184.021.007). This work was carried out on the Dutch national infrastructure with the support of SURF Cooperative.

## Software implementation

R/BioConductor^38,39^ http://bioconductor.org/packages/bacon/

**Methods** see **Supplemental Methods**

**Accession codes**. EGA EGAC00001000277.

#### Inflation factors are variance estimates of the empirical null

Given a set of p z-statistics, *z*_1_,*z*_2_,…, *z_p_* the genomic inflation factor is often calculated as the median of the squared z-statistics divided by 0.456; median 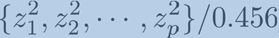, where 0.456 is the theoretical median of a chi-square distribution with one degree of freedom. Alternatively, a “squared” inflation factor can be calculated by instead of squaring the test-statistics taking the absolute value of the test-statistics which leads to test-statistics following a half-normal distribution with theoretical median,0.456^2^ = 0.675. Occasionally, an inflation factor is calculated based on *P* values, i.e., the ratio of the median log10-transformed *P* values with empirical median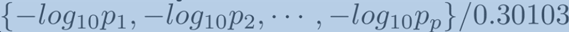 of log10-transformed uniformly distributed statistics (*P* values that follow the null distribution). Actually, median can be used since, 0.30103 is the median of the exponential distribution, log_10_ 2, with rate parameter,log_e_ 10 which is the same as a log10-transformed uniformly distributed random variable.

As a matter of fact, the inflation factor is an estimator of the variance of the z-statistics that follow the null distribution. Therefore, any “robust” estimator of the variance (or standard deviation) could have been used, e.g., the median absolute deviation of the z-statistics (using the appropriate scale factor, 1.4826, for normally distributed statistics).

However, as we show in this paper these approaches are not robust enough when a moderated proportion of true associations is present. A fact that can be proven easily^16^:

“*The median of the squared test-statistics will be the ordered test-statistic at position p/2 or p+1/2, if p is odd or even, respectively. Since, the set of test-statistics represents p_0_ test-statistics following the null distribution and p — p_0_ the alternative with most of them larger than those from the null. The set of ordered test-statistics is roughly given by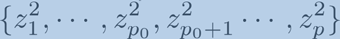. Furthermore, it is known in advance that p_0_ > p/2 or p_0_ > p+1/2, e.g., the proportion of true associations is small, it follows that med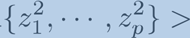 med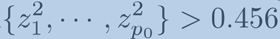 and λ^2^ > 1 even in case no true inflation is present.*”

#### The linear model for association studies

Linear regression is one of the most commonly used models for EWAS/TWAS analysis. In EWAS/TWAS analysis, DNA methylation or gene expression data are considered the dependent variable, in contrast to GWAS where the phenotype of interest is more often considered to be the dependent variable. The linear model for association of the jth feature, i.e., gene or CpG, with the phenotype of interest, **x**, can be given by:

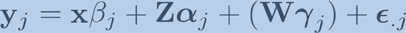

Known technical or biological covariates can simply be added to the model as covariates (**Z**), as well as, estimated unobserved covariates (**W**), e.g., the first few principal components of the data. The corresponding test for association of the j^th^ feature with the phenotype of interest is the regular test for the null hypothesis that the true slope (*β*) of a regression model, is identical to zero.

It has been common, if inflation is present, to divided the test-statistics by the inflation factor before calculation of *P* values. This approach is identical to calculation of *P* values using an inflated or overdispersed normal distribution, with mean zero and variance equal to the squared inflation factor, on the uncorrected test-statistics, i.e, a two-sided association P value is calculated as:

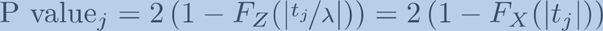

where *Z ~ N (0,1)*, a standard normal distribution and *X ~ N (0,λ^2^)*, an inflated or overdispersed normal distribution.

It has been shown that the omission of covariates not only introduces inflation but bias too^37^, which leads to an empirical null distribution with mean and variance representing bias and inflation. The previous results can easily be translate to fixed-effect meta-analyses. Consider a set of test-statistics *t_ij_* obtained for *i = 1,…, p* features and *j = 1,…, n* cohorts. Bayesian estimates for bias 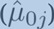 and inflation 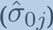 for cohort j are obtained. Then the corresponding bias and inflation corrected test-statistics are given by:

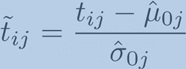

Fixed effect meta-analyses are based on effect sizes and standard errors, since t*_ij_* = *β_ij_/*SE_(*β_ij_*)_, where β_ij_ represents the effect-size and SE(*β_ij_*) the corresponding standard error, we can rewrite the above equation to get corrected effect sizes and standard errors:

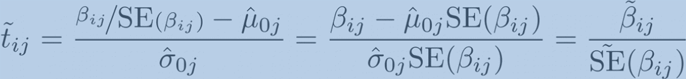

The corrected effect sizes and standard errors can be used in any software to perform fixed effect meta-analysis.

## REFERENCES

1 Rakyan, VK. Down, T.A. Balding, D.J. and Beck, S. Epigenome-wide association studies for common human diseases. Nat. Rev. Genet. 12, 8, (2011).

2 de Magalhaes, J.P. Curado, J. Church, G.M. Meta-analysis of age-related gene expression profiles identifies common signatures of aging. Bioinformatics, 25, 7 (2009).

3 Peters, M.J. et al. The transcriptional landscape of age in human peripheral blood. Nat. Commun., 22, 6 (2015).

4 Hannum, G. et al. Genome-wide methylation profiles reveal quantitative views of human aging rates. Mol Cell. 49, 2 (2013).

5 Beineke P. et al. PREDICT Investigators. A whole blood gene expression-based signature for smoking status. BMC Med Genomics. 5, 58 (2012).

6 Gao X, Jia M, Zhang Y, Breitling LP Brenner H. DNA methylation changes of whole blood cells in response to active smoking exposure in adults: a systematic review of DNA methylation studies. Clin Epigenetics. 7, 113 (2015).

7 Mill, J, Heijmans, B.T. From promises to practical strategies in epigenetic epidemiology. Nat. Rev. Genet. 14, 8, (2013).

8 Devlin, B. and Roeder, K. Genomic control for association studies. Biometrics, 55, 4 (1999).

9 Lehne, B. et al. A coherent approach for analysis of the Illumina HumanMethylation450 BeadChip improves data quality and performance in epigenome-wide association studies. Genome Biology, 15, 16 (2015).

10 Zou, J., Lippert, C. Heckerman, D. Aryee, M. Listgarten, J. Epigenome-wide association studies without the need for cell-type composition. Nat. Meth., 11, 3 (2014).

11 Yang, J. et al. Genomic inflation factors under polygenic inheritance. Eur. J. Hum. Genet., 19, 7 (2011).

12 Voorman, A. Lumley, T. McKnight, B. and Rice, K. Behavior of QQ-plots and genomic control in tudies of gene-environment interaction. PLoS ONE, 6, 5 (2011).

13 Bacanu, S.A. Devlin, B. Roeder, K. Association studies for quantitative traits in structured populations. Genet. Epidemiol., 22, 1 (2002).

14 Devlin, B. Bacanu, S.A. and Roeder, K. Genomic control to the extreme. Nat. Genetics. 36, 11 (2004).

15 Verdinelli, I. and Wasserman, L. Bayesian analysis of outliers problems using the Gibbs sampler. Statistics and Computing. 1 (1991).

16 Efron, B. Large-Scale Simultaneous Hypothesis Testing: The Choice of a Null Hypothesis. JASA, 99, 465 (2004).

17 Efron, B. Size, Power and false discovery rates. The Annals of Statistics. 35, 4 (2007).

18 Schwartzman, A. Empirical null and false discovery rate inference for exponential families. The Annals of Applied Statistic. 2, 4 (2008).

19 Schuemie, M.J. Ryan, P.B. DuMouchel, W. Suchard, M.A. Madigan, D. Interpreting observational studies: why empirical calibration is needed to correct p-values. Stat. Med. 30, 2 (2014).

20 Zhernakova, D. et al. Hypothesis-free identification of modulators of genetic risk factors. http://biorxiv.org/content/early/2015/11/30/033217.

21 Bonder, M.J. et al. Disease variants alter transcription factor levels and methylation of their binding sites. http://biorxiv.org/content/early/2015/11/30/033084..

22 Leek, J.T. and Storey, J.D. Capturing heterogeneity in gene expression studies by surrogate variable analysis. PLoS Genetics, 3, 9 (2007).

23 Gagnon-Bartsch, J.A. and Speed, T.P. Using control genes to correct for unwanted variation in microarray data. Biostatistics. 13, 3 (2012).

24 Wang, J. Zhao, Q. Hastie, T. and Owen, A.B. Confounder Adjustment in Multiple Hypotheses Testing. ArXiv:1508.04178, (2015).

25 Kerr, K.F. Comments on the analysis of unbalanced microarray data. Bioinformatics, 25, 16 (2009).

26 Diebolt, J. and Robert, C.P.Estimation of Finite Mixture Distributions through Bayesian Sampling. JRSS B, 56, 2 (1994).

27 Devlin, B. Roeder, K. Wasserman, L. Genomic control, a new approach to genetic-based association studies. Theor. Popul. Biol., 60, 3 (2001).

28 Leek, J.T. et al. Tackling the widespread and critical impact of batch effects in high-throughput data. Nat. Rev. Genet., 11, 10 (2010).

29 Risso, D. Ngai, J. Speed, T.P. and Dudoit, S. Normalization of RNA-seq data using factor analysis of control genes or samples. Nat. Biotechnol., 32, 9 (2014).

30 Teschendorff, A.E. Zhuang, J. and Widschwendter, M. Independent surrogate variable analysis to deconvolve confounding factors in large-scale microarray profiling studies. Bioinformatics, 27, 11 (2011).

31 Maksimovic, J. Gagnon-Bartsch, J.A. Speed, T.P. and Oshlack, A. Removing unwanted variation in a differential methylation analysis of Illumina HumanMethylation450 array data. Nucleic Acids Res., 43, 16 (2015).

32 Thompson, J.R. Attia, J. and Minelli, C. The meta-analysis of genome-wide association studies. Brief Bioinform. 12, 3 (2011).

33 Anders, S. and Huber, W. Differential expression analysis for sequence count data. Genome Biol. 11, (2010).

34 Robinson, M.D. & Smyth, G.K. Moderated statistical tests for assessing differences in tag abundance. Bioinformatics 23, (2007).

35 Du, P. et al. Comparison of Beta-value and M-value methods for quantifying methylation levels by microarray analysis. BMC Bioinformatics, 11, 587 (2010).

36 Law, C.W.Chen, Y. Shi, W. Smyth, G.K. voom: Precision weights unlock linear model analysis tools for RNA-seq read counts. Genome Biology, 3, 15 (2014).

37 Rao, P. Some notes on misspecification in multiple regression. The American Statistican, 25, 5 (1971).

38 R Core Team. R: A Language and Environment for Statistical Computing. R Foundation for Statistical Computing, Vienna, Austria, http://www.R-project.org/, (2015).

39 Huber, W et al. Orchestrating high-throughput genomic analysis with Bioconductor. Nature Methods, 12 (2015).

